# ACKR3 agonism induces heterodimerization with chemokine receptor CXCR4 and attenuates platelet function

**DOI:** 10.1101/2024.05.10.593491

**Authors:** Valerie Dicenta-Baunach, Zoi Laspa, David Schaale, Manuel Sigle, Alp Bayrak, Tatsiana Castor, Thanigaimalai Pillaiyar, Stefan Laufer, Meinrad Paul Gawaz, Anne-Katrin Rohlfing

**Author notes:** **Corresponding author** Anne-Katrin Rohlfing, PhD, University Hospital Tübingen, Department of Cardiology and Angiology, Eberhard Karls University Tübingen, Otfried-Müller-Str. 10; 72076 Tübingen, Germany.

## Abstract

**Background:** Platelet receptors ACKR3 and CXCR4 play a crucial role in a variety of cardio-vascular diseases. Like most chemokine receptors, CXCR4 is a G protein coupled receptor that induces platelet activation. In contrast, the atypical chemokine receptor 3 (ACKR3) lacks the ability to activate heterotrimeric G proteins and its activation leads to platelet inhibition and attenuates thrombus formation. In nucleated cells, heterodimerization of ACKR3 with CXCR4 regulates CXCL12-dependent signaling. The aim of our study was to investigate the formation of ACKR3/CXCR4 heterodimers in platelets and the subsequent consequences for platelet function.

**Methods and results:** Using a microscopy proximity ligation assay (PLA, Duolink^®^) to screen for CXCR4/ACKR3 heterodimerization inducing compounds, we found that ACKR3 agonism but not conventional platelet agonists or endogen ligands lead to heterodimer formation. To further characterize the formation of ACKR3/CXCR4 heterodimers, we studied the CXCL12-dependent platelet activation via CXCR4. Both, CXCL12-dependent platelet aggregation and collagen-dependent *ex vivo* thrombus formation were significantly downregulated by ACKR3 agonism. Moreover, platelet intracellular calcium and Akt signaling were increased by CXCL12 and again suppressed by ACKR3-specific agonists. Previously, CXCL12 was shown to decrease platelet cAMP levels via CXCR4. Treatment with a specific ACKR3 agonist counteracted this CXCL12/CXCR4-dependent cAMP decrease.

**Conclusion:** Our results reveal that the formation of platelet ACKR3/CXCR4 heterodimers is dependent on ACKR3 rather than CXCR4. Furthermore, ACKR3 agonism induced heterodimerization is associated with mitigating CXCL12/CXCR4-dependent platelet activation possibly by modulating CXCR4-dependent G protein signaling. Our results indicate possible ACKR3 agonist functions and reinforce the potential therapeutic applications of ACKR3 agonists.

Graphical abstract:
CXCR4/ACKR3 heterodimerization in platelets.

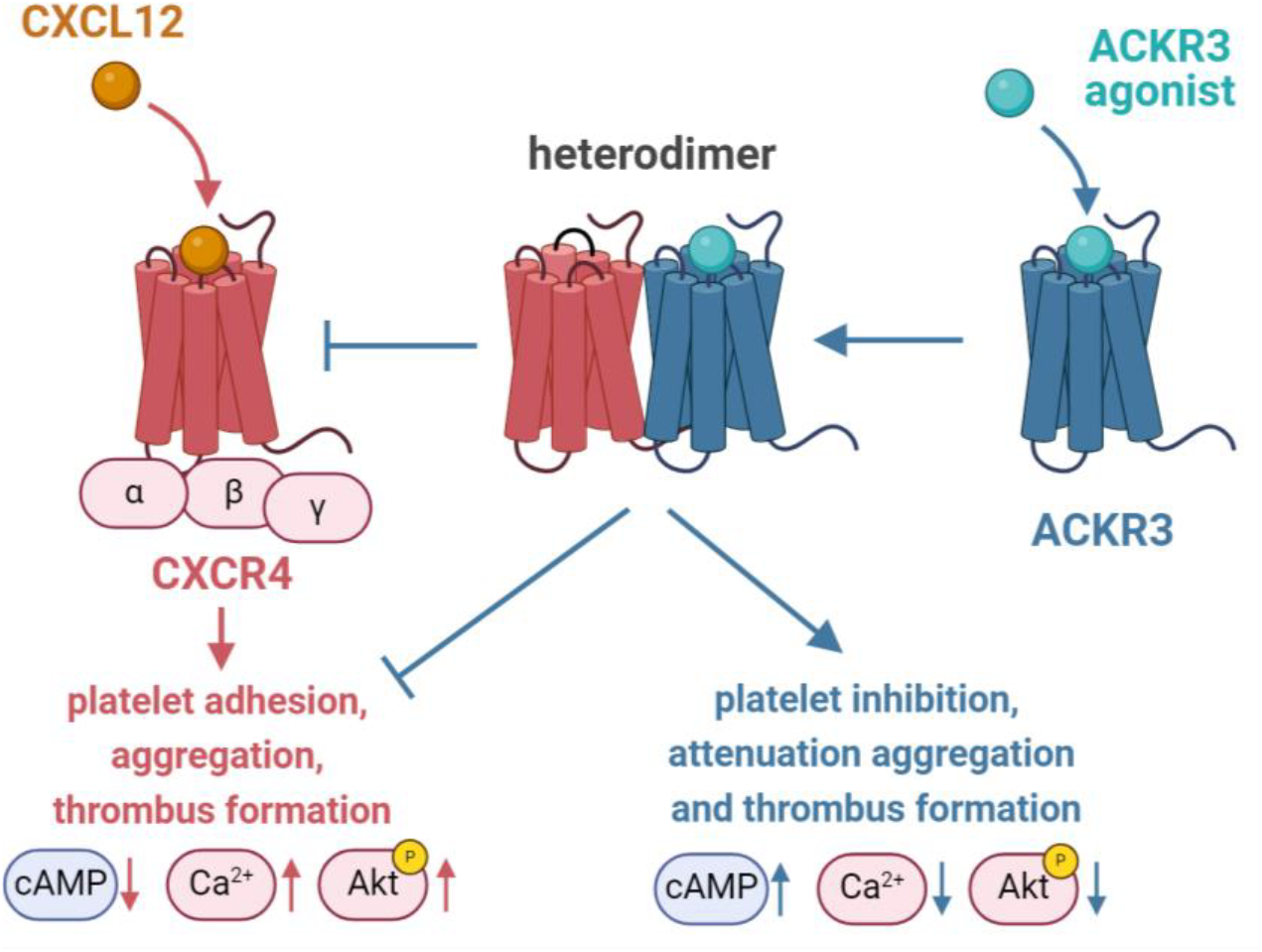

## 1. Introduction

Chemokine receptors play an important role in platelet function.^1-3^ Platelets express a variety of chemokine receptors including CCR1 and CCR4, as well as CXCR4, CXCR6, and CX3CR1.^4-6^ Most chemokine receptors are seven transmembrane-spanning plasma membrane proteins coupled to heterotrimeric G proteins (GPCRs).^7-9^ Interaction of chemokines with their respective chemokine receptors primarily promotes platelet activation.^4-6,10-12^ In contrast, the atypical chemokine receptor 3 (ACKR3, formerly CXCR7) receptor has been recently identified to attenuate platelet activation.^13-16^ Binding of macrophage migration inhibitory factor (MIF) to ACKR3 mitigates activation and prevents platelets from undergoing apoptosis.^15^ Further, genetic deficiency of ACKR3 in mice results in hyperreactivity of platelets^13^ and specific ACKR3 agonists inhibit platelet activation and functions.^13,14^ Surface expression of ACKR3 is dynamically regulated. Stimulation of platelets with CXCL12 (C-X-C motif chemokine ligand 12, also SDF-1α) results in downregulation of CXCR4 and upregulation of ACKR3 on the platelet plasma membrane^16^. This dynamic surface expression of CXCR4 and ACKR3 is regulated via ERK1/2 and cyclophilin A signaling.^16^ In nucleated cells, co-expression of CXCR4 and ACKR3 results in an increased formation of CXCR4/ACKR3 heterodimers and regulates CXCL12-mediated signaling.^17,18^ Whether CXCR4/ACKR3 heterodimerization occurs in anucleated platelets and affects function is unknown.

The purpose of our study was to characterize surface expression and formation of CXCR4/ACKR3 heterodimerization in platelets and to elucidate its consequence for CXCL12-dependent platelet activation and thrombus formation.

## 2. Materials and Methods

### Materials

Recombinant human CXCL12 and CXCL14 were purchased from R&D systems (R&D Systems, Minneapolis, Minnesota, USA). ACKR3 agonist VUF11207 was obtained from Merck (Merck KGaA, Darmstadt, Germany). BY-242 and control compound C46 were manufactured by ourselves. Standard chemicals were purchased from Carl Roth (Karlsruhe, Germany) or Merck (Darmstadt, Germany). Platelet agonist CRP-XL (collagen related peptide) was obtained from CambCol Laboratories (Cambridge, UK), adenosine diphosphate (ADP) from Probe & go Labordiagnostica (Lemgo, Germany) and thrombin from Merck (Merck KGaA, Darmstadt, Germany). Prostaglandin E_1_ (PGE_1_) was purchased from Merck (Merck KGaA, Darmstadt, Germany).

### Preparation of human platelets

Prior blood collection, healthy donors gave informed written consent (local ethics committee vote 141/2018B02). Acid-citrate-dextrose anticoagulated blood was centrifuged at 200 xg for 20 min and the resulting platelet rich plasma was washed in modified Tyrode’s-HEPES buffer (137 mM NaCl, 2.8 mM KCl, 12 mM NaHCO_3_, 5 mM glucose, 0.4 mM Na_2_HPO_4_, 10 mM HEPES, pH 6.5) without bovine serum albumin by centrifuging again at 830 xg for 10 min. After discarding the supernatant, the platelet pellet was resuspended in Tyrode’s-HEPES buffer (pH 7.4) and the platelet count was determined using a SYSMEX cell counter (Sysmex Cooperation, Kobe, Japan).

### Proximity Ligation Assay

The proximity ligation assay Duolink^®^ (Merck KGaA, Darmstadt, Germany) was performed according to the manufacturer’s instructions. 1 x 10^6^ washed platelets in Tyrode’s-HEPES buffer (pH 7.4) supplemented with 1 mM CaCl_2_ were treated as indicated and afterwards fixed with 2% formalin. The non-adherent platelets were centrifuged on coverslips and blocked with 10% donkey serum (Merck KGaA, Darmstadt, Germany) and 1% bovine serum albumin (AppliChem, Darmstadt, Germany) in phosphate-buffered saline. Afterwards, the platelets were incubated at 4°C over night with antibodies against CXCR4 (MAB172, Clone # 44716, R&D Systems, Minneapolis, Minnesota, USA), ACKR3 (ab72100, abcam, Cambridge, United Kingdom) or corresponding IgG controls (Mouse IgG2B Isotype Control MAB004 R&D Systems, Minneapolis, Minnesota, USA and rabbit IgG Isotype Control 3900S Cell Signaling Technology, Danvers, MA, USA). The following PLA steps, binding of the PLA probes (oligonucleotide-labeled secondary antibodies), ligation, and signal amplifications were performed according to the manufacturers protocol. Before mounting, the platelets were stained for 30 min with Phalloidin Alexa Fluor^®^ 488 (ThermoFisher Scientific, Waltham, Massachusetts, USA). Fluorescence microscopy images were taken on a Nikon Eclipse Ti2-A microscope (100x DIC objective, Nikon, Tokyo, Japan). The images were analyzed with the NIS-Elements AR software version 5.21 (Nikon, Tokyo, Japan). The PLA pixel count was evaluated in relation to the phalloidin pixel count and thus the number of platelets.

### Immunoblot Analysis

2.5 x 10^6^ washed platelets in Tyrode’s-HEPES buffer (pH 7.4) supplemented with 1 mM CaCl_2_ were treated as indicated and subsequently lysed using RIPA lysis buffer. The protein amount was measured using a standard Bradford assay. Platelet lysates were electrophoretically separated by sodium dodecyl sulfate-polyacrylamide gel electrophoresis (SDS-PAGE, 10%) under reducing conditions and transferred onto a polyvinylidene difluoride membrane. Membranes were blocked with Roti-Block (Carl Roth, Karlsruhe, Germany) or 3% bovine serum albumin (Merck KGaA, Darmstadt, Germany) in TRIS-TWEEN-buffered saline and incubated overnight with the primary antibodies as indicated. Antibodies against phospho-Akt and total Akt were obtained from Cell Signaling Technology (Danvers, MA, USA) and loading control anti-GAPDH was purchased from ThermoFisher Scientific (Waltham, Massachusetts, USA). IRDye secondary antibodies (Li-Cor, Lincoln, Nebraska, USA) were used in a 1:15,000 dilution in Roti-Block or 3% bovine serum albumin/ TRIS-TWEEN-buffered saline. For fluorescence detection and analysis, the LI-COR Odyssey System (Li-Cor, Lincoln, Nebraska, USA) was used.

### Platelet Aggregometry

Platelet aggregation was measured with citrate-anticoagulated platelet rich plasma (1 x 10^8^ platelets/ sample) using a light transmission aggregometer (Aggregometer 490-X; Chrono-Log Corp., Havertown, Pennsylvania, USA)^19^. Platelets were treated as indicated and aggregation was measured for 5 min with a stir speed of 1000 rpm at 37°C. The extent of aggregation was quantified in percentage of light transmission and analyzed using Aggrolink8 software (Chrono-Log Corp., Havertown, Pennsylvania, USA).

### Ex vivo thrombus formation

For *ex vivo* thrombus formation experiments, a flow chamber system (Maastricht Instruments B. V., Maastricht, The Netherlands) with collagen-coated cover slips (100 µg/ml Kollagen Reagens HORM Suspension, Takeda, Tokyo, Japan) at a shear rate of 1,000 s^-1^ was used. Washed platelets (5 x 10^5^ in Tyrode’s-HEPES buffer (pH 7.4) were diluted 4:5 in phosphate-buffered saline containing 1.2 mM Ca^2+^. Subsequently, platelets were treated as indicated and stained for 10 min with fluorochrome 3,3’-dihexy-loxacarbocyanine iodide (DiOC_6_, Sigma Aldrich, St. Louis, Missouri, USA). For visualization, a Nikon fluorescence microscope was used (Nikon Eclipse Ti2-A, 20x objective, Nikon, Tokyo, Japan) and at least 5 images of independently selected areas were taken after the perfusion. The images were analyzed with the NIS-Elements AR software version 5.21 (Nikon, Tokyo, Japan).

### Calcium Signaling

Platelet intracellular Ca^2+^ was measured as described previously using 5 µM Fluo-4 fluorescence dye (Invitrogen, Waltham, Massachusetts, USA).^20^ 1 x 10^5^ washed platelets were labeled with Fluo-4 for 30 min at room temperature (RT), incubated on fibrinogen (100 µg/ml, Sigma Aldrich, St. Louis, Missouri, USA) coated coverslips for 20 min and pretreated with or without 100 µM ACKR3 agonists or control C46 for 15 min at RT. After incubation, non-adherent platelets were removed, fresh Tyrode’s-HEPES buffer (pH 7.4) was added and platelets were analyzed using a fluorescence microscope (Nikon Eclipse Ti2 A; DIC 100x oil objective, Nikon, Tokyo, Japan). Fluo-4 fluorescence was recorded for 90 s and after 15 s platelets were activated as indicated. To estimate the cytosolic Ca^2+^ activity, the fold change in the mean intensity of single platelet fluorescence was determined using the NIS-Elements AR software version 5.21 (Nikon, Tokyo, Japan).

### Measurement of cAMP levels

To determine platelet levels of cyclic adenosine monophosphate (cAMP) an enzyme-linked immunosorbent assay (ELISA, Enzo Life Sciences, Farmingdale, New York, USA) was used. Platelets were treated as indicated and lysed using a lysis buffer containing 0.5% TritonX-100 and 1 mM IBMX (3-Isobutyl-1-methyl-Xanthin) to stop endogenous phosphodiesterase activity. After centrifugation at 12,000 rpm for 5 minutes at RT, the supernatant was used to perform the ELISA according to the manufacturer’s instructions.

### Statistical analysis and graphical presentation

Data are provided as means ± SD and n represents the number of biological replicates. Repeated measures of one-way ANOVA or mixed-effects analysis with an appropriate post-test, as indicated, were performed for multiple comparisons. Two-sided Student’s t-tests were utilized for two group comparison (p <0.05 statistical significance, 95% confidence interval) or Wilcoxon matched-pairs signed rank test for not normally distributed data. All statistical analyzes were performed with GraphPad Prism (GraphPad Software, Inc., La Jolla, CA, USA, Version 10.1.1). Schematic drawings were created using BioRender.com.

## 3. Results

### Heterodimers of ACKR3 with CXCR4 occurs on the platelets surface upon ACXCR3 agonism

Platelets dynamically surface express both ACKR3 and CXCR4 upon activation and in response to CXCL12.^16,21^ To explore heterodimerization of both receptors we developed a Duolink^®^ proximity ligation assay (PLA) using a combination of specific anti-ACKR3 and anti-CXCR4 antibodies (**Figure 1A**). PLA allows quantifying steric interaction of membrane receptors using oligonucleotide-labelled secondary antibodies.^22,23^ Platelets were stimulated with agonists such as collagen-related peptide (1 µg/ml CRP-XL), adenosine diphosphate (5 µM ADP) or thrombin (1 U/ml). Upon activation of platelets with these classical agonists, we did not observe CXCR4/ACKR3 heterodimerization (**Figure 1B**). Previously, we demonstrated that the CXCR4 chemokine CXCL12 decreases expression levels of CXCR4 and enhances surface exposure of ACKR3.^16^ Thus, we asked whether CXCL12 or CXCR4 ligand CXCL14 induce heterodimerization of both chemokine receptors. As shown in **Figure 1C/D** CXCL12 and CXCL14 did not stimulate CXCR4/ACKR3 heterodimerization. Next, we asked, whether ACKR3 agonists promote interaction of CXCR4 with ACKR3. Interestingly we found that specific ACKR3-selective agonists (VUF11207, BY-242) significantly enhanced formation of CXCR4/ACKR3 heterodimers on the platelet surface (p<0.05) (**Figure 1C/D**). Thus, we conclude that heterodimer formation between ACKR3 and CXCR4 occurs on the platelet surface specifically following ACKR3 agonism.

**Figure 1.**
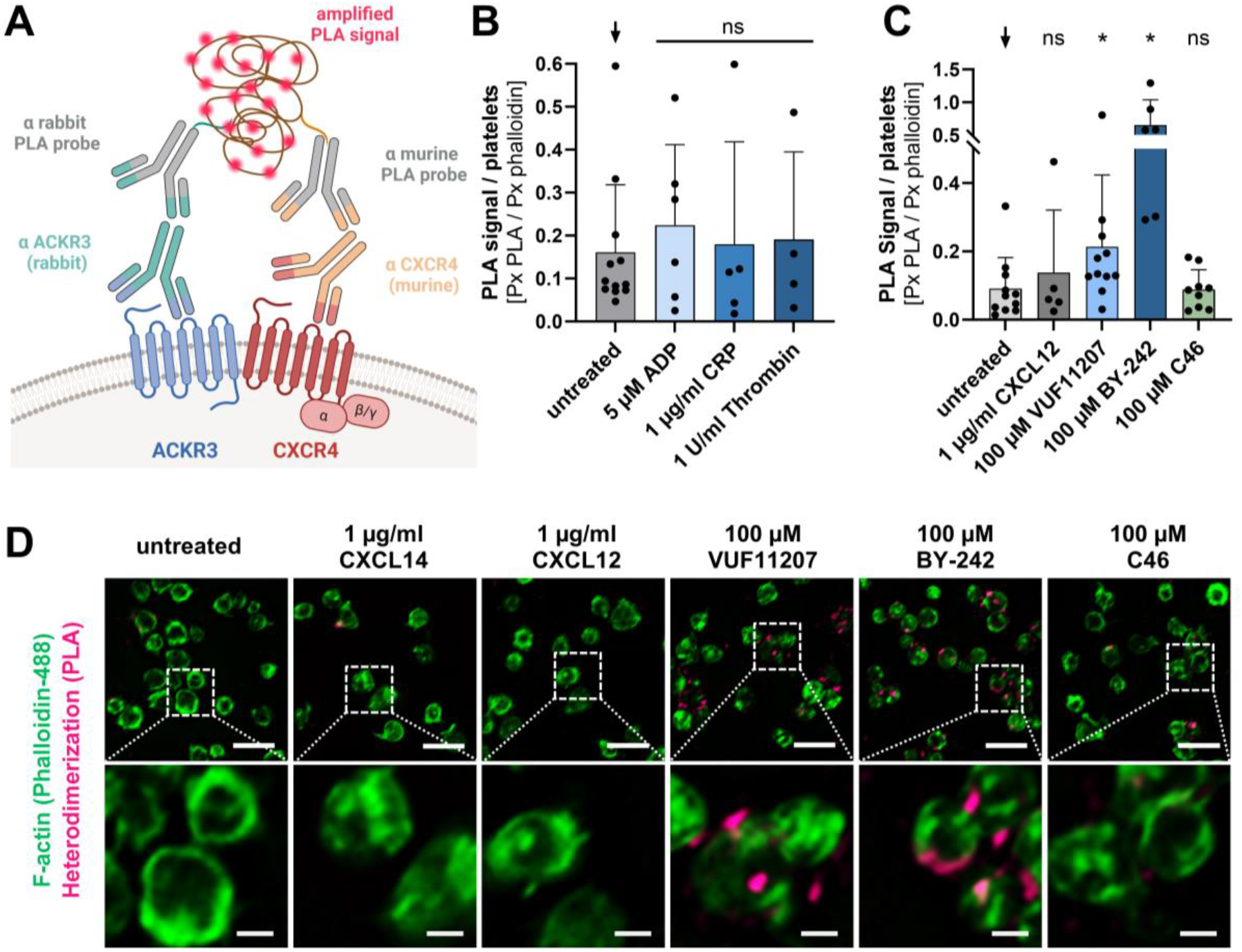
Heterodimers of ACKR3 and CXCR4 occur on platelets surface upon ACXCR3 agonism. **A** Scheme of proximity ligation assay Duolink^®^ functioning. ACKR3 and CXCR4 are labeled with specific primary antibodies of different species. These are detected with the corresponding PLA probes with short reverse DNA tails, which are linked in the ligation step if the two receptors are in close proximity to each other. The signal is amplified by a polymerization reaction. Created with BioRender.com. **B/C** Statistical analysis of PLA for ACKR3-CXCR4 interaction after **B** platelet activation for 30 min with 5 µM ADP, 1 ug/ml CRP-XL and 1 U/ml thrombin compared to untreated control and **C** platelet treatment with 1 µg/ml CXCL12, 1 µg/ml CXCL14 or 100 µM ACKR3 agonist (VUF11207, BY-242) and 100 µM control substance C46 for 15 min at RT. **B/C** Plotted: Arithmetic means ± SD of PLA signal per platelet signal, n≥4, Wilcoxon matched-pairs signed rank test; n.s. not significant, ^**\***^ p < 0.05. **D** Representative images of PLA (magenta) and phalloidin (green) staining of untreated platelets or treated platelets with 1 µg/ml CXCL12, 1 µg/ml CXCL14, or 100 µM ACKR3 agonist VUF11207 and BY-242 and control substance C46 for 15 min. Upper panel: scale bar = 5 µm, lower panel: scale bar = 1 µm.

### ACKR3 agonists attenuate CXCL12-dependent platelet aggregation and thrombus formation under flow

CXCL12 induces platelet activation and aggregation via ligation of CXCL12 with CXCR4.^4,12,24^ In our present study, we confirm that CXCL12 stimulates platelet aggregation in a dose-dependent manner (**Figure 2A**). To test whether ACKR3-agonist-induced CXCR4/ACKR3 receptor heterodimerization is associated with platelet function, isolated human platelets were stimulated with recombinant CXCL12 (1 µg/ml) and the platelet response was analyzed by light aggregometry in the presence and absence of ACKR3 agonists (**Figure 2B**). We found that in the presence of ACKR3 agonists (VUF11207, BY-242) but not a control chemical (C46), CXCL12-induced platelet aggregation was significantly inhibited (control vs. VUF11207: p < 0.05 and control vs. BY-242: p < 0.05) (**Figure 2B**). Next, we tested the effect of ACKR3 agonism on CXCL12-dependent platelet thrombus formation on immobilized collagen under flow. As described for aggregation, CXCL12 increased thrombus formation (untreated vs. CXCL12: p < 0.05). Platelet-dependent thrombus formation in response to CXCL12 was substantially reduced (p < 0.01) in the presence of ACKR3 agonists (VUF11207, BY-242) but not the control chemical C46 (**Figure 2C**). Our data imply that ACKR3 agonist-dependent CXCR4/ ACKR3 heterodimerization is associated with a reduction of platelet function in response to CXCL12.

**Figure 2.**
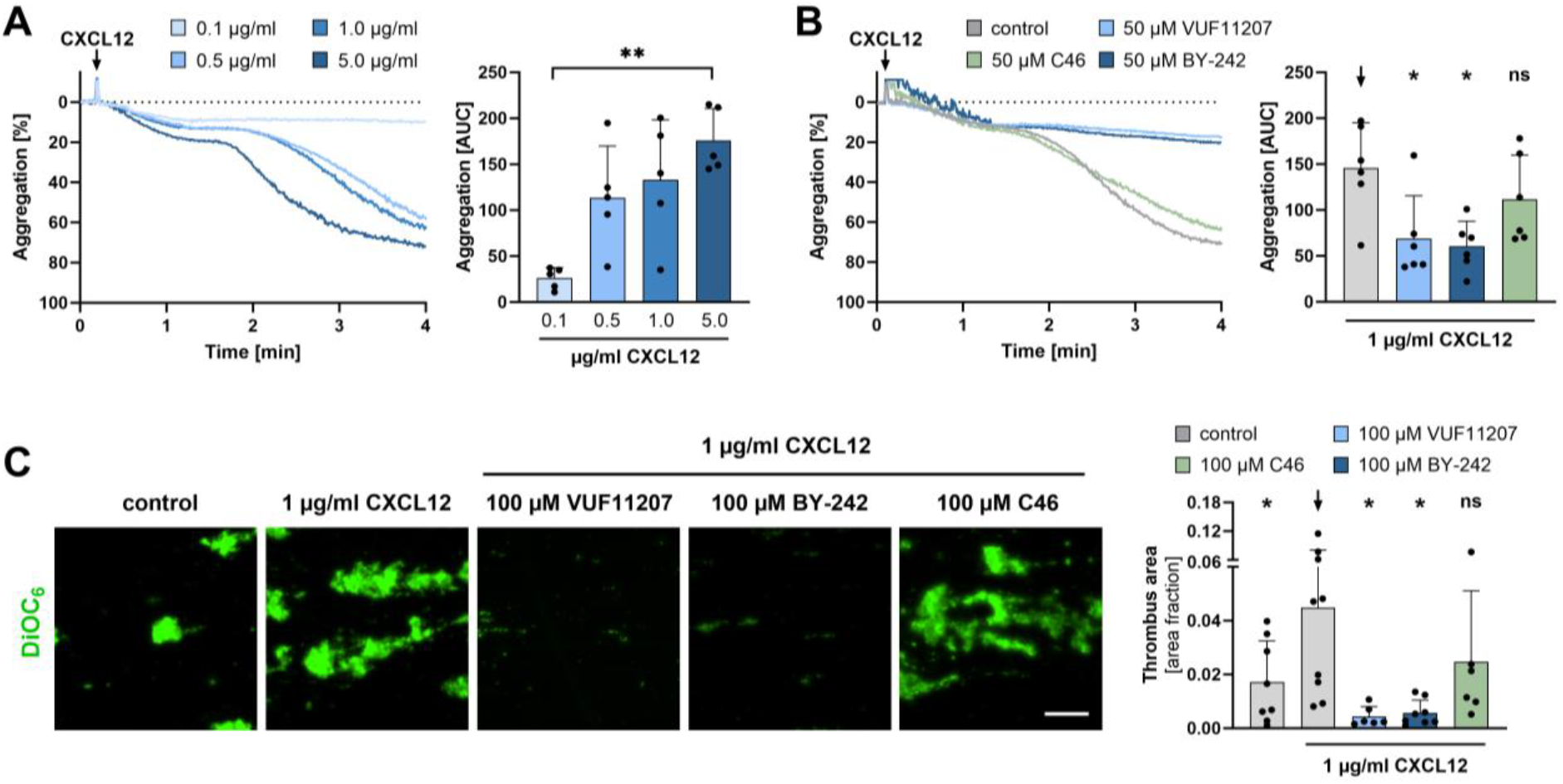
ACKR3 agonist-induced heterodimerization of ACKR3/CXCR4 is associated with attenuation of CXCL12-dependent platelet function. **A** Representative light transmission aggregation curves (left) and statistical analysis (right) of platelets treated with 0.1, 0.5, 1.0, and 5.0 µg/ml CXCL12. Plotted: Arithmetic means ± SD of platelet aggregation, n=5, ordinary one-way ANOVA; not indicated = n.s. not significant, ^**\*\***^ p < 0.01. **B** Representative light transmission aggregation curves (left) and statistical analysis (right) of platelets pretreated with 50 µM ACKR3 agonist (VUF11207, BY-242) or 50 µM control substance C46 for 15 min at 37°C and activated with 1.0 µg/ml CXCL12. Plotted: Arithmetic means ± SD of platelet aggregation, n=6, ordinary one-way ANOVA; not indicated = n.s. not significant, ^**\***^ p < 0.05. **C** Representative microscope images of DiOC_6_ stained platelets (left) and statistical analysis (right) of *ex vivo* thrombus formation. Washed platelets were pretreated with 100 µM ACKR3 agonist (VUF11207, BY-242) or 100 µM control substance C46 for 15 min at RT and activated with 1.0 µg/ml CXCL12 for 10 min. Plotted: Arithmetic means ± SD of thrombus area fraction, n≥6, Wilcoxon matched-pairs signed rank test or student’s t-test; n.s. not significant, ^**\***^ p < 0.05.

### CXCL12-dependent platelet signaling is inhibited by ACKR3 agonism

Recently, we described that ACKR3 ligation modulates platelet activation signaling.^13,25^ To test the effect of ACKR3 agonism and CXCL12-dependent Ca^2+^ signaling, platelets were loaded with the fluorescent calcium indicator Fluo-4 and treated with CXCL12 (1 µg/ml) (**Figure 3A**). CXCL12 substantially induced an increase in intracellular Ca^2+^. This CXCL12-induced Ca^2+^ signaling was significantly attenuated in the presence of ACKR3 agonist compared to controls (p<0.001) (**Figure 3A**). Further, CXCL12 induced Akt phosphorylation in a concentration-dependent manner (**Figure 3B**). As described above for intracellular Ca^2+^ signaling, CXCL12-dependent Akt phosphorylation was significantly reduced in the presence of ACKR3 agonist VUF11207 (CXCL12 vs. CXCL12 + VUF11207: p < 0.01) (**Figure 3C**). Previously, CXCL12 has been shown to initiate platelet thrombus formation through CXCR4 and changes in cAMP and Ca^2+^ signaling^12^. Furthermore, cAMP has been shown to inhibit Akt phosphorylation and thereby AKT-mediated signaling.^26^ Thus, we asked whether ACKR3 agonism counteracts CXCL12-dependent cAMP suppression. Indeed, we detected that CXCL12-dependent suppression of cAMP (p < 0.05) is partially reversed in the presence of ACKR3 agonist (CXCL12 vs. CXCL12 + VUF11207: p < 0.05) (**Figure 3D**).

**Figure 3.**
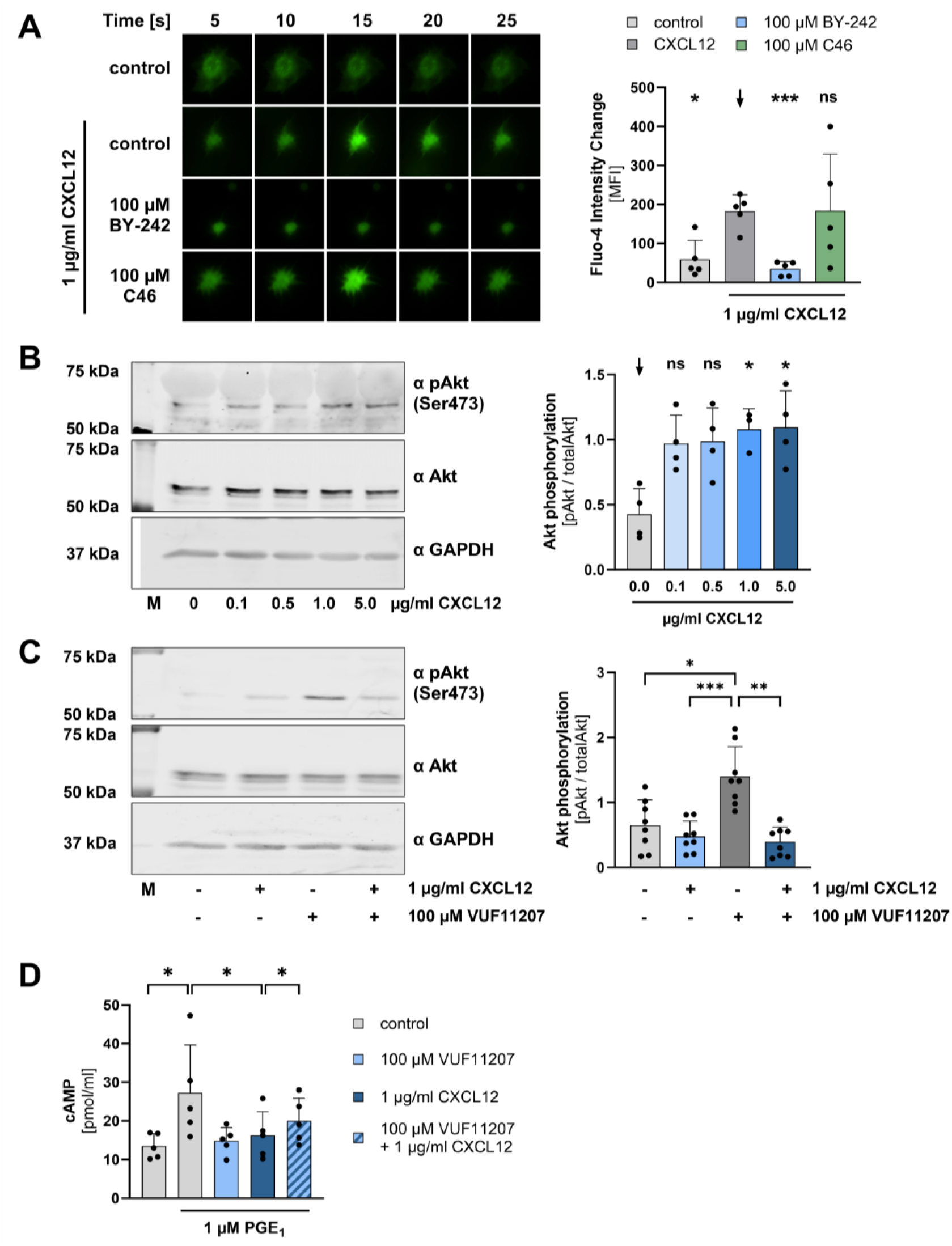
CXCL12-dependent platelet signaling is inhibited by ACKR3 agonism. **A** Intracellular calcium signaling of platelets using Fluo-4 staining (30 min RT). Representative microscope images (left) and statistical analysis (right) of washed platelets pretreated with 100 µM ACKR3 agonist BY-242 or 100 µM control substance C46 for 15 min at RT and activated with 1.0 µg/ml CXCL12. Plotted: Arithmetic means ± SD of Fluo-4 intensity change, n=5, ordinary one-way ANOVA; n.s. not significant, ^**\***^ p < 0.05, ^**\*\*\***^ p < 0.001. **B** Representative images of platelets Akt phosphorylation and statistical analysis of washed platelets treated with 0 – 5.0 µg/ml CXCL12 for 15 min at RT. M = Protein size marker. Plotted: Arithmetic means ± SD of fold-change of pAkt/Akt, n≥3, Mixed-effects analysis Dunnett’s multiple comparisons test; n.s. not significant, ^**\***^ p < 0.05, ^**\*\***^ p < 0.01. **C** Representative images of platelets Akt phosphorylation and statistical analysis of washed platelets pretreated with or without 100 µM ACKR3 agonist VUF11207 and subsequent platelet activation with 1 µg/ml CXCL12 compared to untreated control. M = Protein size marker. Plotted: Arithmetic means ± SD of fold-change of pAkt/Akt, n=8, ordinary one-way ANOVA Dunnett’s multiple comparisons test; ^**\***^ p < 0.05, ^**\*\***^ p < 0.01, ^**\*\*\***^ p < 0.001. **D** cAMP levels of washed platelets pretreated with or without 100 µM ACKR3 agonist VUF11207 for 15 min at RT and subsequent platelet treatment with 1 µM PGE_1_ and 1 µg/ml CXCL12 for 10 min at 37°C. Plotted: Arithmetic means ± SD of cAMP levels, n=5, student’s t-test; not indicated = n.s. not significant, ^**\***^ p < 0.05.

In summary, ACKR3-induced heterodimerization of CXCR4/ACKR3 is associated with significant inhibition of platelet aggregation and thrombus formation. A function that is regulated via Ca^2+^-cAMP-Akt signaling (**schematic drawing, Figure 4**). Our data imply that heterodimerization of ACKR3 with CXCR4 modulates the responsiveness of CXCR4 towards its ligand CXCL12.

**Figure 4.**
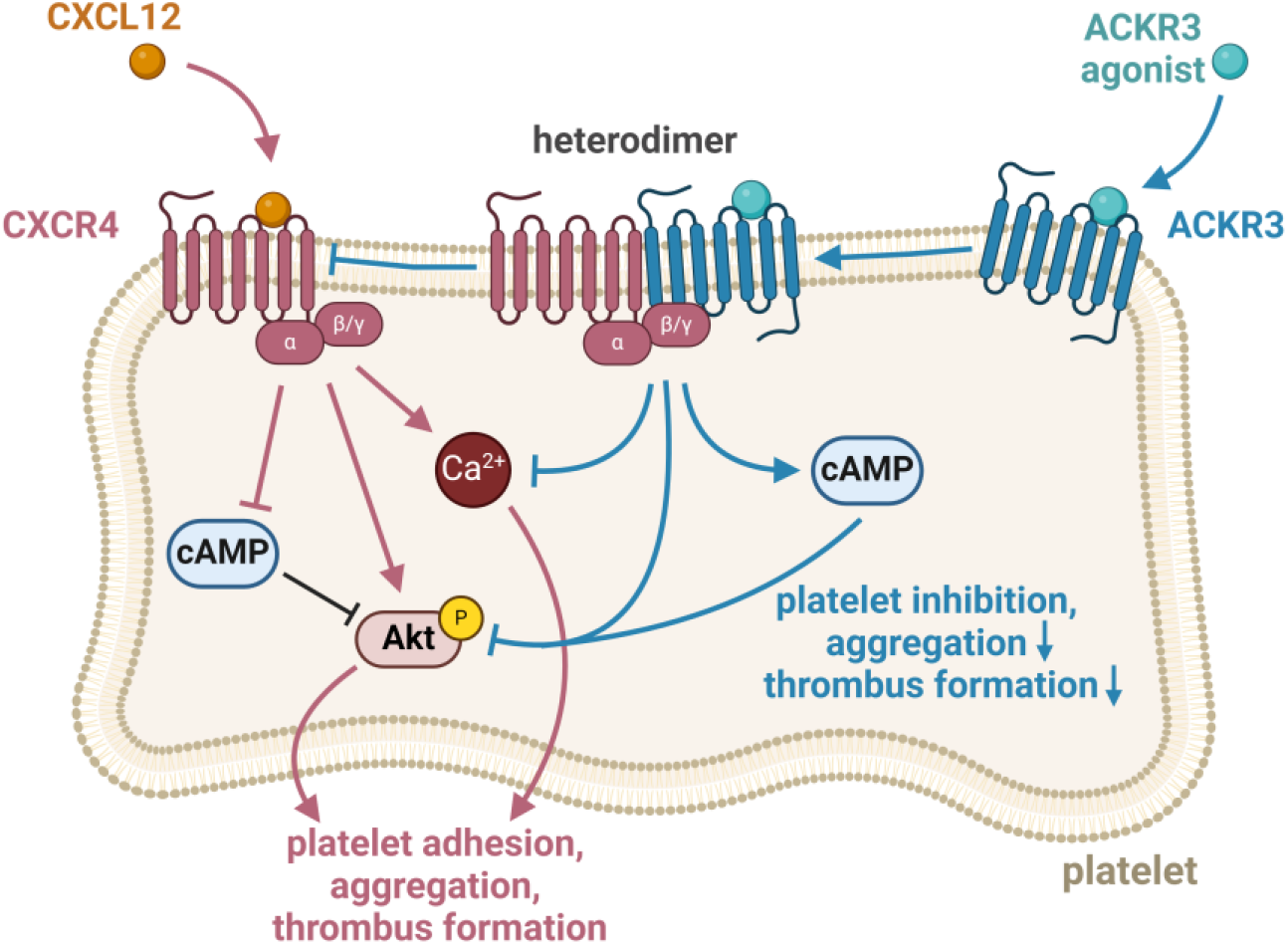
CXCR4/ACKR3 heterodimer associated signaling in platelets. Schematic drawing of CXCR12 dependent-signaling via CXCR4. ACKR3 agonism induced CXCR4/ACKR3 heterodimerization mitigates CXCL12/CXCR4-dependent platelet adhesion, aggregation, and thrombus formation by counteracting cAMP inhibition and attenuation of Akt and Ca^2+^ signaling. Created with BioRender.com

## 4. Discussion

The major findings of the present study are the following: i) Selective ACKR3 agonism induces ACKR3 heterodimerization with CXCR4 ii) ACKR3 agonists attenuate CXCL12-dependent platelet aggregation and thrombus formation. iii) CXCL12-dependent platelet signaling is mitigated by ACKR3 agonism. Unlike most other chemokine receptors, ACKR3 is a platelet inhibitory receptor.^13,15^ The endogenous CXCR4 and ACKR3 ligand MIF inhibits platelet activation and apoptosis via ACKR3.^15^ Recently, it has been shown that patients suffering from coronary artery disease with elevated ACKR3 levels had an improved clinical prognosis and platelets showed less aggregation.^13,25^ The murine platelet-specific ACKR3 knockout exhibits hyperreactive platelets, leading to increased *ex vivo* thrombus formation and platelet activation as well as degranulation.^13^ In addition, animals carrying this platelet specific *Ackr3* knockout showed increased damage and inflammation of the myocardium and brain after ischemia/reperfusion.^13^ Targeting ACKR3 has been shown to inhibit platelet activation.^14^ Previously, the use of ACKR3 agonists has been shown to inhibit human and murine platelet activation and aggregation and to increase cAMP levels and the amount of antiplatelet lipids.^25^ In addition, in murine *in vivo* experiments, agonists were able to reduce thrombo-inflammation and infarct size without affecting basal hemostasis.^13,25^ Those findings encouraged us to investigate the role of ACKR3 agonism in CXCR4/ACKR3 heterodimerization and platelet function. In the present study, we show that the formation of platelet CXCR4/ACKR3 heterodimers is dependent on ACKR3 agonism. ACKR3 agonism, but not platelet agonists (such as ADP or collagen-related peptide) nor the CXCR4 ligands CXCL12 and CXCL14 increased significantly the PLA signal of CXCR4/ACKR3 heterodimers. To get further insights in the role of CXCR4/ACKR3 heterodimers for platelets, we investigated CXCL12-dependent platelet function and signaling. CXCL12 is the endogenous ligand of ACKR3 and CXCR4, whereby the binding to the latter in platelets leads to internalization and subsequent ACKR3 externalization.^16^ CXCL12 is secreted, inter alia, by platelets and is therefore a paracrine and autocrine platelet agonist^10,27^ especially potentiating other platelet agonists such as ADP and fibrinogen.^4,28^ These findings are in line with our aggregation and thrombus formation experiments. CXCL12 induced concentration-dependent platelet aggregation and increased the thrombus area onto immobilized collagen under flow. Both, CXCL12-dependent aggregation and thrombus size were significantly attenuated using specific ACKR3 agonists. Previously, CXCL12 was shown to induce platelet aggregation via CXCR4 rather than ACKR3.^12,24^ Together with our findings, this indicates that CXCR4/ACKR3 heterodimerization is associated with the inhibition of CXCL12-dependent platelet activation via CXCR4. Unlike ACKR3, which lacks the ability to activate heterotrimeric G proteins, CXCR4 is a G protein coupled receptor (GPCR). Analog to the ADP receptor P2Y_12_, CXCR4 is coupled to G_αi_ and CXCL12 ligation leads to G_αi_-dependent inhibition of the adenylyl cyclase (AC).^10,12^ Thus, the level of platelet modulating cAMP is decreased. Indeed, PGE_1_-induced platelet cAMP levels were reduced using CXCL12 and again, ACKR3 agonism counteracted this effect. Further, CXCL12-dependent Akt and calcium signaling was diminished using a selective ACKR3 agonist. Lou *et al*. demonstrated that cAMP is able to inhibit Akt phosphorylation.^26^ Our data indicate that AC-cAMP signaling is modulated by CXCL12 ligation to CXCR4. Furthermore, by CXCL12-dependent reduction of the cAMP levels, Akt phosphorylation and intracellular calcium are increased, which is counteracted by ACKR3 agonism. Previously, it was shown that heterodimerization of CXCR4 and ACKR3 in HEK293T cells changed the ability of CXCR4 to interact with its G proteins.^18^ Thus, it is conceivable that ACKR3 agonism leads to CXCR4/ACKR3 heterodimerization, which is associated with the modulation of CXCL12-dependent interaction of CXCR4 with its coupled heterotrimeric G proteins. Platelet ACKR3 and CXCR4 play a crucial role in a variety of cardio vascular diseases^13,21,29^ and it was recently shown that platelet-specific CXCL12 knockout mice also show limited arterial thrombosis without prolonging the bleeding time.^30^ Thus, this expands the possible use of specific ACKR3 agonists for the therapeutic regulation of platelets.

## Abbreviations

ACKR3: atypical chemokine receptor 3 (formerly CXCR7)
ADP: adenosine diphosphate
CRP-XL: collagen related peptide
CXCL12: C-X-C motif chemokine ligand 12
CXCR4: C-X-C motif chemokine receptor type 4
DiOC_6_: 3,3’-dihexyloxacarbocyanine iodide
GPCR: G protein coupled receptor
MIF: macrophage migration inhibitory factor
PGE_1_: prostaglandin E_1_
PLA: proximity ligation assay

## Acknowledgements / Grants

This project was supported by the Deutsche Forschungsgemeinschaft (DFG, German Research Foundation) – Project number 335549539 – GRK2381. D.S. was supported by a research grant of the Deutsche Gesellschaft für Kardiologie-Herz und Kreislaufforschung (DGK, German Cardiac Society).

## Data Availability

For original data please contact.

## Authors’ contributions

V.D.-B., M.P.G. & A.-K.R. conceived and designed research. V.D.-B., Z.L., D.S., M.S., A.B., performed experiments & analyzed data. V.D.-B., Z.L., D.S., M.S., M.P.G. & A.-K.R. interpreted results of experiments. V.D.-B., M.P.G. & A.-K.R. prepared figures and drafted manuscript. Z.L., D.S., M.S., A.B., T.C., T.P., S.L., M.P.G., A.-K.R. edited and revised manuscript. All authors have read and agreed to the published version of the manuscript.

